# The *Caenorhabditis elegans* RAD-51 isoform A is required for the induction of DNA-damage-dependent apoptosis

**DOI:** 10.1101/152058

**Authors:** Marcello Germoglio, Adele Adamo

**Affiliations:** Institute of Biosciences and BioResources, National Research Council, via Pietro Castellino 111, 80131 Naples, Italy; University of Campania “Luigi Vanvitelli”, Viale Abramo Lincoln, 5, 81100 Caserta Italy

**Keywords:** apoptosis, genome instability, homologous recombination, meiosis, C. elegans

## Abstract

The *rad-51* gene in *Caenorhabditis elegans* is transcribed into alternative mRNAs potentially coding three alternative protein isoforms. We have genetically modified this gene in order to investigate the potential roles of the longest isoform, namely isoform A, in genome stability. The RAD-51 isoform A appears to contribute to genome stability in late development, but is not implicated in meiosis or DNA repair in the germline. However, the RAD-51 isoform A has a pivotal role in DNA damage induced apoptosis, but not in DNA damage checkpoint activation or physiological cell death. This is a relevant new finding that improves our understanding of how DNA damage apoptosis is restricted to late pachytene stage preventing the inappropriate loss of nuclei undergoing the earlier stages of meiotic recombination, during which a large number of physiologically induced DSBs are present.

## Introduction

Homologous repair is the most faithful double strand break (DSB) repair program. However, it can only occur when a homologous substrate is present: either the sister chromatid after replication or the homologous chromosome in meiosis. Single strand invasion is the essential step in initiating such a process. The key protein at this step is the RecA-like protein Rad51. This protein is conserved in all eukaryotes from fungi to humans (La Volpe and Barchi 2012).

Meiosis is the mechanism that ensures proper segregation of homologous chromosomes during eukaryotic gametogenesis in order to confer a complete haploid genome to oocytes and sperms (Kleckner 1996; Hillers *et al.* 2017). During meiosis the topoisomerase-like protein Spo11 physiologically induces DSBs. Homologous repair will then occurs among homologous chromosomes in order to generate cross-overs (COs) and subsequent chiasmata that are essential for generating appropriate tension between homologous chromosomes and faithful segregation at anaphase I (Kleckner 1996).

In most eukaryotes a meiosis specific RecA-like protein, Dmc1, is required in the strand invasion step during homologous recombination at meiosis (Bishop *et al*. 1992). Rad51 and Dmc1 cooperate in this process (Brown and Bishop 2014). Surprisingly, in some organisms, such as in Sordariomycetes, *Caenorhabditis elegans,* and *Drosophila melanogaster*, the key actor in the meiotic step of strand invasion, Dmc1, is missing and Rad51 alone seems to perform strand-invasion in meiosis.

The *C. elegans* germ line exhibits the complete time course of meiotic prophase I in which nuclei at the different stages of oogenesis can be easily identified based on their position and chromosome appearance. In wild-type *C. elegans,* RAD-51 foci, representing nascent meiotic DSBs, arise during zygotene and early pachytene stages and then decrease in number during the middle pachytene stage as meiotic DSBs repair progresses (Colaiacovo *et al.* 2003).

Increase and/or persistence of such foci are indicative of DNA repair defects. During early meiotic prophase there is a clear bias towards homologous recombination using the homologous chromosome as a template, thus promoting CO formation (Rosu *et al.* 2011). In order to perform COs, chromosomes must be aligned and stabilized by the synaptonemal complex (SC) (Colaiacovo *et al.* 2003). As in other organisms, there is a surveillance system in *C. elegans* that checks recombination intermediates and pairing formation during meiosis (Machovina *et al.* 2016). During meiotic prophase, a number of proteins cooperate to ensure CO formation: some are involved in promoting pairing and synapsis of homologous chromosomes (Couteau and Zetka 2005; Colaiacovo 2006; Penkner *et al.* 2007; Meneely *et al.* 2012), whereas others, known as pro-CO factors, seem to be involved in stabilizing recombination intermediates between homologous chromosomes (Zalevsky *et al.* 1999; Kelly *et al.* 2000). In absence of the SC, as in *syp-2* mutants, COs do not occur. Later, residual DSBs can be repaired using the sister chromatide. At this stage DSBs repair is *brc-1* dependent (Adamo *et al.* 2008).

By the end of pachytene, about half of the oocytes in the *C. elegans* germ line undergo physiological cell death (Gumienny *et al.* 1999). In addition, exogenously induced DNA damage or persistent DNA lesions, due to endogenous DNA repair defects, are sufficient to trigger the DNA-damage checkpoint and significantly increase germline apoptosis (Gartner *et al.* 2000; Schumacher *et al.* 2005; Adamo *et al.* 2008; 2010; Rutkowski *et al.* 2011). In addiction to the DNA damage checkpoint, a different checkpoint enhances apoptosis during pachytene in response to defects in SC assembly (Bhalla and Dernburg 2005).

When exogenous DNA damage occurs, the DNA damage checkpoint induces cell cycle arrest at the G2 phase in the premeiotic proliferative region of the germ line (Schumacher 2001 *et al.*; Moser *et al.* 2009) to allow DNA repair to occur. The DNA damage checkpoint requires the 9-1-1 complex, a DNA damage–sensor complex that contains the conserved proteins RAD-9, RAD-1, and HUS-1 (Hofmann *et al.* 2002). Many proteins are involved in the activation of such sensor complex such as, for example, the ZTF-8 protein directly interacting with RAD-1. In order to fulfill this role, ZTF-8 needs to be SUMOylated (Kim and Colaiacovo 2014, 2015). Following 9-1-1 loading, activation of the p63-like protein CEP-1 is required to elicit apoptosis (Schumacher *et al.* 2001). Important to note that germline apoptosis only occurs in nuclei in the late pachytene stage of meiosis suggesting that there must be specific signals that make late pachytene nuclei competent for apoptosis and that prevent apoptosis at earlier meiotic stages. Restricting DNA damage-induced apoptosis to late pachytene stage could prevent the accidental loss of nuclei undergoing the first steps of meiotic recombination, during which DSBs are present (Colaiacovo *et al.* 2003, Adamo *et al.* 2012). Some genes, whose action is essential for apoptosis induction in presence of DNA damages during *C. elegans* gametogenesis, appear to act downstream of the proapoptotic signaling (Silva *et al.* 2013, Ackermann *et al.* 2016). Some of these genes are also involved in CO formation such as *msh-4, msh-5,* and *zhp-3* (Silva *et al.* 2013, Jantsch *et al.* 2004). Another gene, involved in DNA damage induced apoptosis, is the *ufd-2* gene coding for the E4 ubiquitin ligase UFD-2. In its absence, RAD-51 foci persist after pachytene, and DNA damage-induced apoptosis is prevented (Ackermann *et al.* 2016). Interestingly, nuclei undergoing apoptosis in late pachytene appear overloaded of the RAD-51 protein although DNA repair would be useless at this stage (Colaiacovo *et al.* 2003).

By the diakinesis stage, chromosomes are highly structured and appear as six DAPI stained bodies per nucleus. Each body corresponds to a couple of homologous chromosomes held together by chiasmata that are the result of the occurred COs. In CO defective mutants, such as *syp-2*, we can observe a vast majority of diakinesis nuclei containing twelve DAPI stained bodies corresponding to the 12 univalent chromosomes.

*C. elegans rad-51* gene inactivation leads to a severe phenotype: unstructured chromosomes at diakinesis, reduced brood-size, high levels of germline apoptosis, and 100% embryonic lethality at the second generation (Gartner *et al.* 2000; Rinaldo *et al.* 2002; Alpi *et al.* 2003).

When the *C. elegans rad-51* gene was first isolated (Rinaldo *et al.* 1998), it was shown that the *C. elegans* RAD-51 is positioned in a phylogenetic tree (derived by bootstrap analysis) at an intermediate branch between the eukaryotic Rad5l group and that of the eukaryotic Dmc1. The Rad5l fungal sequences surprisingly seem to score much better than the *C. elegans* RAD-51 sequence against the vertebrate Rad51. Although the *C. elegans* RAD-51 protein is more similar to members of the Rad51 family than to the Dmc1 family, some conserved Dmc1-specific amino acids are retained suggesting that the *C. elegans* RAD-51 may represent a hybrid performing the activities of both these strand-invading proteins. Therefore, in *C. elegans* a single gene locus is responsible for all the functions ascribed to the two loci in fungi as well as in vertebrates. The *rad-51* genomic locus, composed of eight exons, was shown to be transcribed into two alternative messanger RNA (Rinaldo *et al.* 1998). More recent evidences suggest the presence of a third intermediate transcript. The second exon is only present in the longer transcript. At the time of the isolation of the gene, the authors hypothesized that the two different isoforms coded by this gene might be used at different stages and/or in different tissues (Rinaldo *et al.* 1998). The extra amino acids at the amino-terminal end of the long isoform might confer a different tissue-specificity or function to the coded protein. We here investigate this possibility.

## Materials and Methods

### Strains

*C. elegans* strains (listed in Supplemental Material, Table S1) were cultured at 20°C on NGM plates with *Escherichia coli* OP50 as food source according to standard methods (Brenner 1974).

### PCR primers

The PCR to identify *rad-51(A^-^)* mutation are performed with three primers:

Upper *rad-51*: 5’TATGGGACAATCTTGGGG3’

Upper *rad-51(A^-^)*: 5’ACGAAGTTATGGCTGCTCG3’

Lower *rad-51*: 5’ACGAAGTTATGGCTGCTCG3’

The PCR to identify *brc-1(tm1145)* mutation are performed with two primers:

Upper *brc-1*: 5’TGTCGCATCGTCGGCATTAA3’

*Lower brc-1:* 5’AATATAGGCACCGGCGGGGA3’

### Screening of laying worms

Worms were isolated and cloned during the L4 larval state on Petri plates and left at 20°C to lay eggs for 3 days. They were transfered every 12 hours to lay eggs onto fresh plates until the deposition of non-fertilized oocytes. Each plate content was monitored every for 24/72 hours to analyze the following parameters among the progeny: embryonic lethality, presence of males, aberrant phenotypes and larval arrests. The percent of embryonic lethality is calculated as the ratio of unviable eggs on laid eggs. The percentage of males, aberrant phenotypes and larval arrests is calculated as the ratio of males/aberrant/larval phenotype on the hatched eggs.

Number of scored P_0_ worms: *wt*: 20; *rad-51(A^-^*): 12; *cep-1*: 8.

### Larval growth assay

To analyze the larval growth rate after IR exposure, nematodes were synchronized with Alkaline Hypochlorite solution 1:1 (NaOH 1.6M, NaClO 0.6%) and embryos were transferred on seeded plates. Following the hatching of eggs the L1 worms are exposed to 120 Gy γ-rays, using a Caesium-137 source and observed after 53 hrs to analyze two different parameters: larval growth rate and the frequency of aberrant phenotypes.

Number of scored worms: *wt* 0 Gy: 105; *rad-51(A^-^)* 0 Gy: 114; *wt* 120 Gy: 97; *rad-51(A*^-^) 120 Gy:65

### Immunostaining of meiotic nuclei

Gonads from 20 hrs post L4 adults were dissected in M9 solution (0.3% H2PO4, 0.6% Na2HPO4, 0.5% NaCl and 1mM MgSO4). Slides were frozen in liquid nitrogen, then immersed at −20°C in methanol, methanol/acetone (1:1) and acetone respectively for 5 min, and washed three times in PBS for 5 min each time. Slides were left in 3% BSA in PBS for 30 min at 37°C in a humid chamber. Primary rabbit antibody α-RAD-51 was diluted 1:200 in Ab buffer (1% BSA, 0.1% Tween-20, 0.05% sodium azide in 1X PBS). Slides were incubated for 90 min at room temperature followed by three washes in PBS for 5min. Secondary antibody Texas red anti-rabbit was diluted 1:400 in Ab buffer. Slides were incubated for 60 min in darkroom at room temperature, followed by three washes in PBS + 0.1% Tween-20 for 5 min. Slides were mounted with Prolong Gold Antifade reagent with DAPI (Life Technologies).

To perform the immunostaining after IR exposure, young adult worms were transferred on seeded plates and exposed to 120 Gy γ-rays, using a Caesium-137 source. Following the treatment, worms were incubated for 60 min at 20°C before the gonads dissection. The quantitative analyses of RAD-51 foci were performed by dividing the germ line into 6 zones (gonadal tip, mitotic zone, transition zone, early pachytene, middle pachytene and late pachytene stage), in accordance with their cytological features.

Number of scored nuclei are indicated in the legends

The quantifications of nuclei positive for RAD-51 1hr, 16hrs and 48hrs after IR were performed by immunostaining after 20 Gy γ-rays exposure of young adult worms, using a Caesium-137. Following the treatment, worms were incubated for 1hr, 16hrs and 48hrs at 20°C before the gonads dissection.

Number of scored nuclei are indicated in Table 2.

### Quantitative analysis of germline apoptosis

Adult nematodes were suspended in M9 solution and stained by incubation with 33 μM SYTO-12 (Molecular probes) for 1hr and 30 min at room temperature in the dark. The worms were then transferred to seeded plates to allow stained bacteria to be purged from the gut. After 30 minutes, the animals were mounted on 2% agarose pads in 2mM levamisole. The estimation of apoptotic levels for each genotype was calculated as the average number of apoptotic nuclei per gonadal arm.

Number of scored arms are indicated in the legends.

### Analysis of DAPI-stained germ lines

Adult nematodes were suspended in M9 solution on glass slides, permeabilized and fixed with 10 μl of 100% Et-OH. Finally, slides were mounted in 10 μl of DAPI (2ng/μl) diluted in M9. Number of scored nuclei are indicated in the legends of charts.

Mitotic nuclei were scored 15 h after 120 Gy γ-rays exposure of young adult worms, using a Caesium-137 Number of observed nuclei: *wt* 0 Gy: 85, *rad-51(A^-^)* 0 Gy:130; *wt* 75 Gy: 72, *rad-51(A^-^)* 75 Gy: 106.

### Image collection and processing

Collection of images was performed using a Leica DM6 fluorescence microscope, Hamamatsu camera under the control of Leica LAS AF 6000 software. Images were processed using Leica LAS AF 6000 software and Image J program. Quantitative analysis of RAD-51 foci and DAPI-stained bodies along the germline were performed on z series, Optical sections were collected at 0.18 μm and 0.50 μm increments respectively.

### Statistical tools

Statistical analyses of apoptosis levels and RAD-51 foci patterns were computed through t-Student test for independent samples.

The level comparison of aberrant phenotypes and larval arrests of the different genotypes were computed through χ ^2^ test.

Statistical analyses of nuclear diameters at the premeiotic tip were computed throught the two-tailed Mann-Whitney U test.

### Statement on data availability

Strains are available upon request.

File S1 contains the Table Figures:

Table S1 (list of all strains used in this study and description *rad-51(A^-^)* mutant CRISP/Cas9-mediated genome editing); Table S2 (Statistical analysis of RAD-51 foci in meiosis: t-student); Table S3 (Statistical comparing of apoptotic nuclei in various genetic combination: t-student)

File S2 contains the Supplemental Figures:

Figure S1 (*rad-51(A^-^)* mutant by CRISP/Cas9 mediated genome editing); Figure S2 (RAD-51 A in the larval development); Figure S3 (genetic cross); Figure S4 (Immunostaining of RAD-51 in *syp-2; rad-51(A^-^)* strains).

## Results

### Lack of the RAD-51 isoform A affects genome stability in late development stages, but does not alter COs formation in Meiosis I

In order to understand whether the different isoforms coded by the *rad-51* gene perform different functions in *C. elegans,* we edited the *rad-51* gene by mean of the CRISPR Cas9 technology obtaining a mutant that cannot express the long RAD-51 isoform A (see Material and Methods and figure S1), but retains the ability to code for the shorter forms. We called the mutant *rad-51(A^-^).*

We observed the presence of occasional larval arrests in the mutant population, but we did not observe an increase in the level of males or dead embryos that would be suggestive of chromosome non disjunction in meiosis (Table 1). We then synchronized the nematodes and exposed the larvae at the stage L1/L2 to 120 Gy of g-rays and screened for developmental defects after 52 hrs. We observed in the *rad-51(A^-^)* mutant a significant increase in the larval arrest, eggless, and vulvaless phenotypes compared to the wild type in the same conditions (Table 2 and Figure S2) suggesting a requirement of the RAD-51 isoform A in maintainance of genome stability during late development.

We further questioned the efficiency of CO and chiasma formation in our mutant by cytogenetic analysis: *rad-51(A^-^)* shows six DAPI stained bodies (corresponding to the wildtype six bivalents) in all the diakinesis nuclei (Figure 1A). Furthermore, we assayed the frequency and distribution of RAD-51 foci along the adult hermaphrodite gonad by mean of immunolocalization and we observed no significant differences from the wild type strains (Figure 1B and Table S2). RAD-51 foci start to appear on chromosomes in the transition zone, increase in early pachytene and are unloaded in middle to late pachytene, as in the wild type. Therefore, CO formation does not seem to be affected by the lack of RAD-51 isoform A.

**Figure 1.**
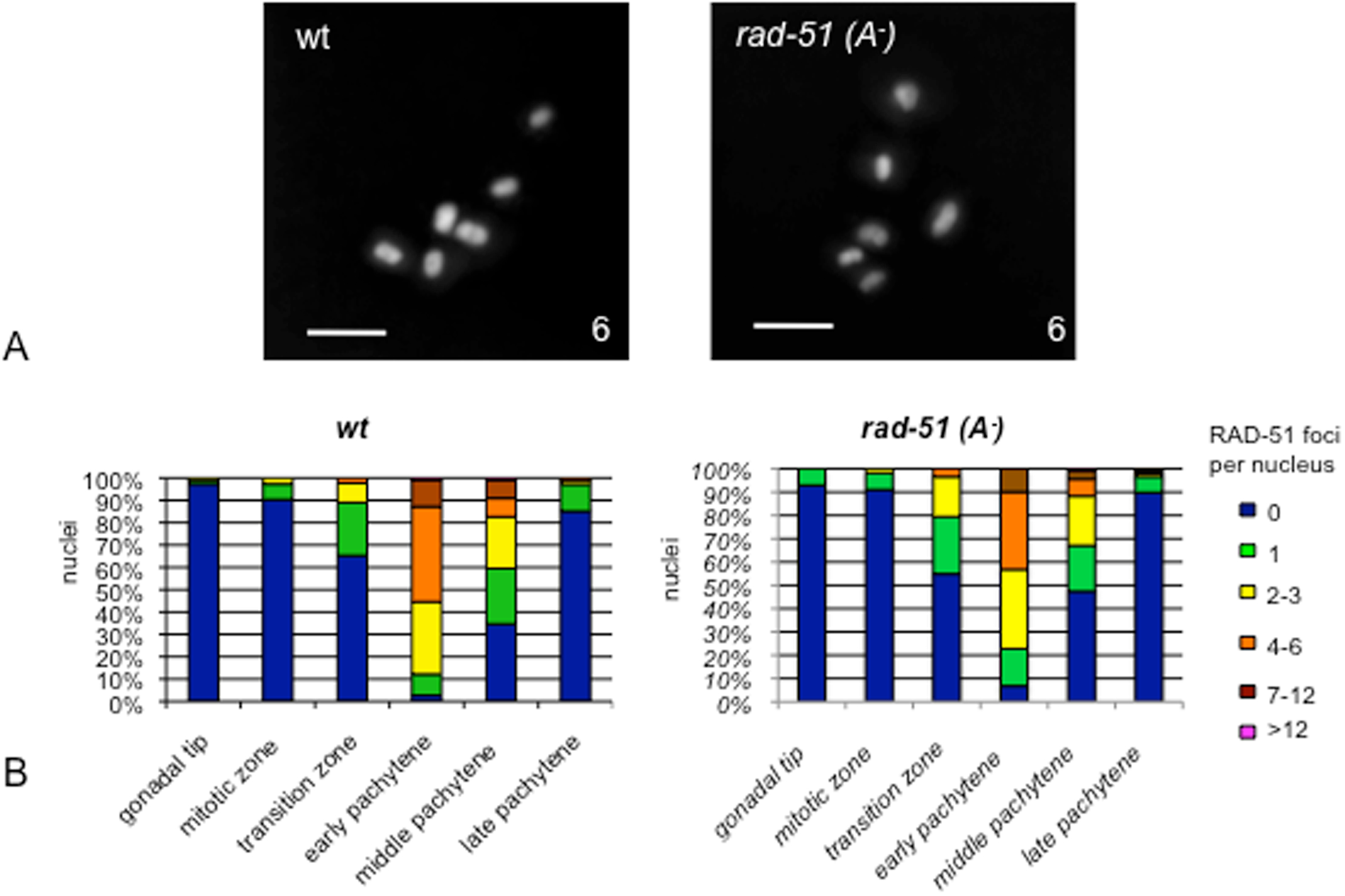
*rad-51(A^-^)* mutant shows six DAPI stained bodies in diakinesis nuclei and exibits a frequency and distribution of RAD-51 foci similar to the wild type. (A) Representative images of diakinesis nuclei of the indicated genotype stained with DAPI. The number of DAPI-stained bodies is indicated at the bottom right of each panel. Number of observed diakinesis nuclei: *wt=* 95, *rad-51(A^-^)=* 92. Scale bar, 2 μm. (B) Quantification of RAD-51 foci along the germlines of the indicated genotypes. An average of 160 nuclei for each gonad region were scored for *wt* and *rad-51(A^-^)*. Statistical analyses are reported in table S2.

### RAD-51 isoform A is not involved in inter-sister DNA repair during meiosis

In wildtype *C. elegans,* one and one only CO per pair of homologous chromosomes is observed. However, as in other eukaryotes, each homologous pair is subject to more than one DSB performed by SPO-11. It has been reported that meiotic DSBs are about twice the number of the final COs in *C. elegans* (Mets and Meyer 2009).

Some DNA repair genes such as *brc-1* and *fcd-2* are thought to be involved in meiosis in the intersister homologous repair of those DSBs that have not been resolved as COs (Adamo *et al.* 2008; 2010). We wondered whether RAD-51 isoform A might be loaded at such a step and might be involved in such a pathway. It has been demonstrated that, in absence of the synaptonemal complex, BRC-1 is necessary to repair the DSBs that could have not been resolved as COs: in fact, while in the *syp-2* single mutant about 80% of diakinesis nuclei show 12 DAPI stained bodies corresponding to the 12 univalents, in the double mutant *syp-2; brc-1* only a third of the nuclei have 12 DAPI stained bodies, while the large majority shows fragmented chromosomes at diakinesis (Adamo *et al.* 2008). We therefore assayed the number of diakinesis DAPI stained bodies in the double mutant *syp-2; rad-51(A^-^).* In this background most diakinesis nuclei show 12 DAPI stained bodies (Figure 2A) and a very small proportion (about 15%) shows chromosome fragmentation similarly to what observed in the *syp-2; cep-1* germline (Figure 2B).

**Figure 2.**
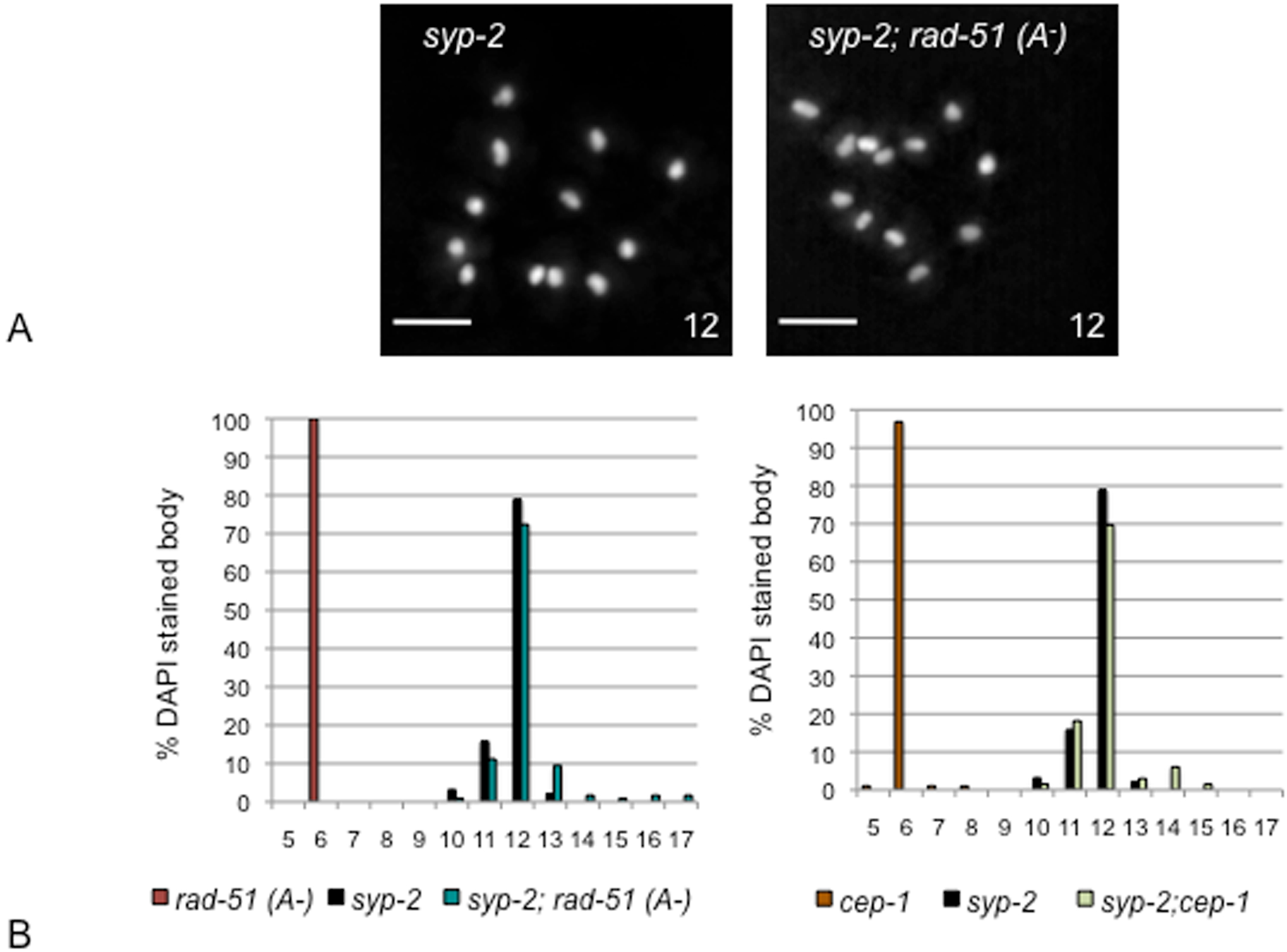
*rad-51(A^-^)* mutant is able to performe inter-sister homologous repair. A) Representative images of diakinesis oocytes of the indicated genotype stained with DAPI. The number of DAPI-stained bodies is indicated at the bottom right of each panel. Scale bar 2 μm. (B) Histograms represent quantification of the DAPI-stained bodies. The number (n) of observed nuclei is indicated next to each genotype. The y axis represents the percentage of nuclei in each class and the x axis indicates the number of DAPI-stained bodies. Genotypes are indicated in the color legend at the bottom of the chart.

We can conclude that *rad-51(A^-^)* mutant is able to accomplish COs formation and inter-sister homologous repair during meiosis.

### RAD-51 isoform A is necessary for DNA damage apoptosis in response to exogenously induced DSBs

In order to assay the sensitivity to exogenous damages in the *rad-51(A^-^)*mutant, we treated it with 120 Gy of γ-rays along with the wild type and the *cep-1* strains. As observed for other phenotypes after γ-ray treatment, in the *rad-51(A^-^)* mutant RAD-51 foci appear at very high level from the mitotic tip of the gonad up to the end of the pachytene stage (Figure 3). However, unlike what occurs in wild type, in *51(A^-^).* as well as in *cep-1,* no increase in apoptosis levels, compared to untreated nematodes, was observed (Figure 4A and Table S3). A possible explanation for this phenomenon is that the long isoform A, coded by the *rad-51* gene, may be necessary to activate the DNA damage dependent apoptosis pathway. However, this is in contrast with what has been observed so far in experiments on gonads in which all the isoforms of the RAD-51 protein were absent. RNA-interference-mediated gene inactivation of the *rad-51* locus, in fact, has been shown to induce, even in absence of gamma rays, a high level of DNA-damage-dependent apoptosis that can be suppressed by the *cep-1* mutation (Gartner *et al*. 2000). Similarly, the *rad-51(lg8701)* mutant strain, carrying a large deletion of the locus and therefore defective for all the isoforms, also shows a high level of apoptosis (Alpi *et al.* 2003; Silva *et al.* 2013). We therefore tried to explore this contradiction. The *rad-51(lg8701)* mutant in homozygousis produce a 100% lethal progeny, therefore only the F1 progeny of heterozygous parents can be studied. We wondered if a relative small amount of maternal RAD-51 isoforms might be present in the adult gonad. We therefore performed a RAD-51 immunolocalization on the *rad-51(lg8701)* F1 progeny of heterozygous parents and detected a very low, but significant amount of RAD-51 foci in the gonad (Figure 4B). We hypothesised that the relative small amount of maternal RAD-51 isoforms B and C might be insufficient to repair the DNA damages, however the traces of RAD-51 isoform A may still allow the damage detection and the activation of the DNA damage dependent apoptosis pathway. In order to demonstrate that this small amount of maternal protein may be necessary and sufficient to activate the DNA damage dependent apoptosis, we assayed the apoptosis of the *rad-5 (lg8701)* homozygous progeny of *rad-51 (lg8701)*;*rad-51(A^-^)* heterozygous hermaphrodite parents (see figure S3 for details of the genetic cross). These nematodes could have inherited the RAD-51 short isoforms B and C by maternal effect, but could not have inherited the long isoform A that has not been expressed for two generations. These nematodes, unlike what observed so far either in the *rad-51*RNAi or in the *rad-51(lg8701)* knockout strain at the F1, only show (low?)basal level of apoptosis supporting our model (Figure 4C and Table S3).

**Figure 3.**
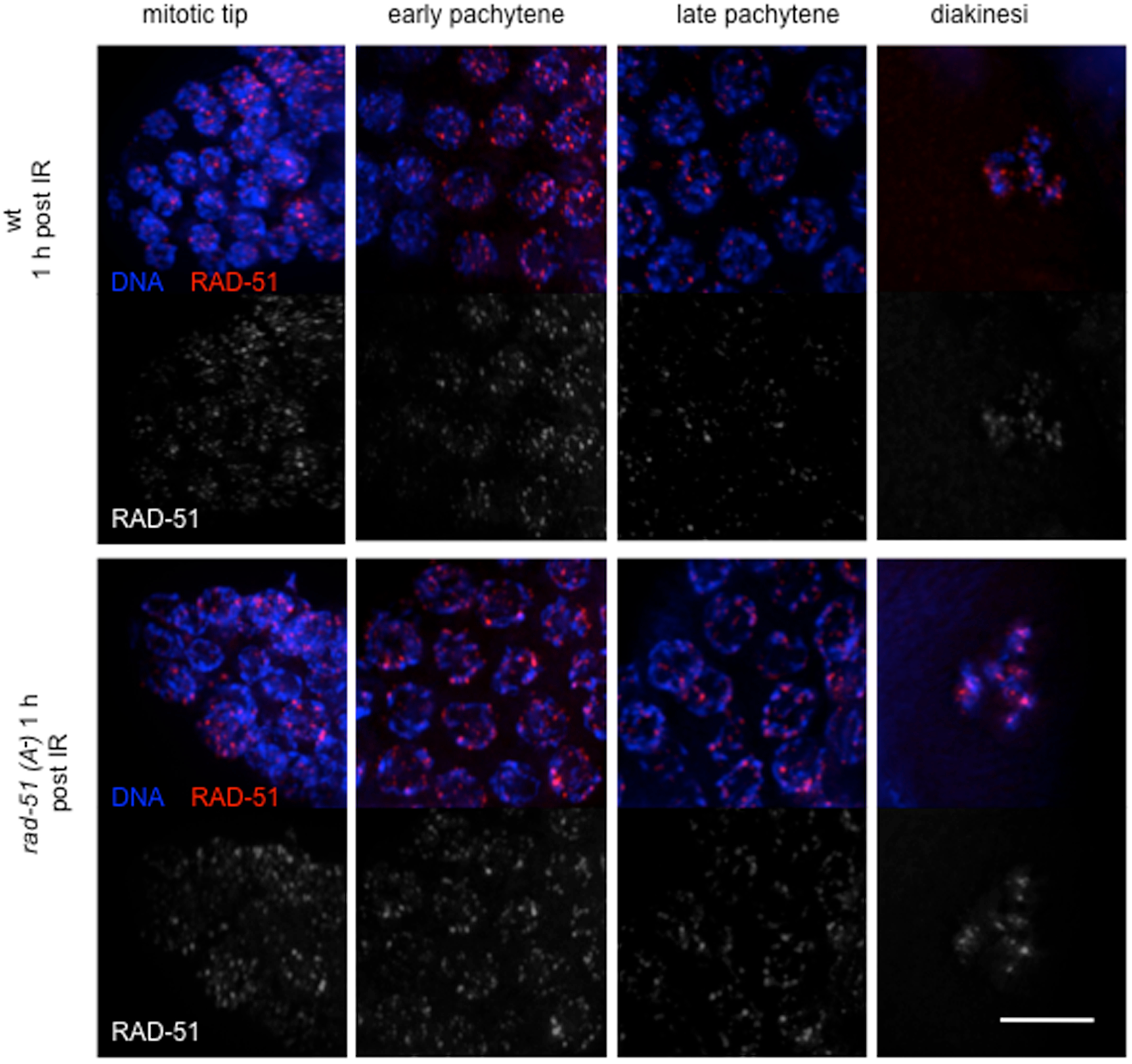
RAD-51 foci in different regions of the gonad after 1hour post IR. Representative images of indicated regions of the gonads stained with anti-RAD-51 antibodies (in red) and with DAPI (in blue) in wt and *rad-51(A^-^)* strains 1 hour post irradiation (120 Gy). Scale bar, 5 μm.

**Figure 4.**
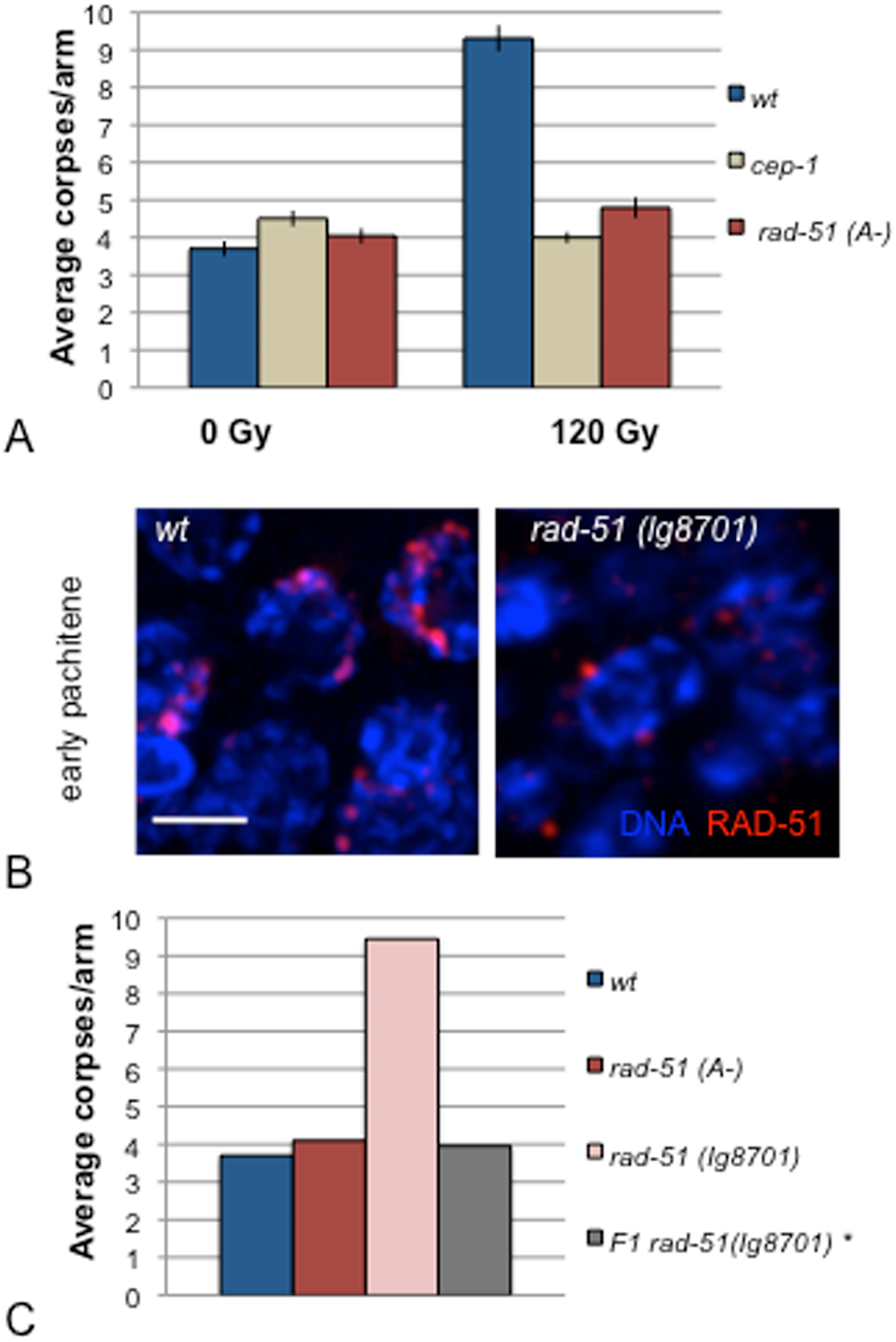
RAD-51 A is necessary to induce germ-nuclei apoptosis after irradiation. (A) Average apoptosis levels in the indicated genotypes. The y axis shows the average number of SYTO-12 labeled nuclei per gonadal arm. Number of observed gonads: *wt* 0 Gy: 71, *cep-1* 0 Gy: 82, *rad-51(A^-^)* 0 Gy: 88, *wt* 120 Gy: 61, *cep-1* 120 Gy: 68, *rad-51(A^-^)* 120 Gy: 69. Error bars correspond to standard error of the means (S.E.M.) calculated from at least three independent experiments. Student’s *t*-tests are shown in table S3. (B) Immunolocalization with anti-RAD-51 in early pachytene nuclei of the *rad-51(lg8701)* mutant shows a low but detectable number of foci (in red). Chromosomes were stained with DAPI (in blue). Scale bar, 5 μm. (C) Average apoptosis levels in the indicated genotypes. The y axis shows the average number of SYTO-12 labeled nuclei per gonadal arm *rad-51(lg8701)*\* represents the *rad-51(lg8701)* homozygous progeny of *rad-51(lg8701)*;*rad-51(A^-^)* heterozygous hermaphrodites. Number of observed gonads: *wt* : 71, *rad-51(A^-^)* : 88, *rad-51(lg8701):21, rad-51 (lg8701)*\*: 24. Student’s *t*-tests are shown in table S3.

### Ionizing radiation triggers the DNA damage checkpoint in the mitotic proliferative region of the *rad-51(A^-^)*gonad

The nuclei in the distal tip of the *C. elegans* gonad are engaged in mitotic proliferation. They respond to exogenous DNA damage triggering a temporary block in cell cycle progression to allow DNA repair. As a consequence of the cell cycle arrest, nuclei in the mitotic region of the germ line of irradiated worms appear enlarged and their overall number is reduced. In order to understand if the RAD-51 isoform A is involved in checkpoint activation or in apoptosis induction, we measured the nuclei diameters in the mitotic region before and after irradiation at 75 Gy. As shown in figure 5A, the *rad-51(A^-^)* mutant is proficient in activating the DNA damage response following IR. In fact, the average diameter of mitotic nuclei increases to twice their original size as is also observed in the wild-type control (Figure 5B).

**Figure 5.**
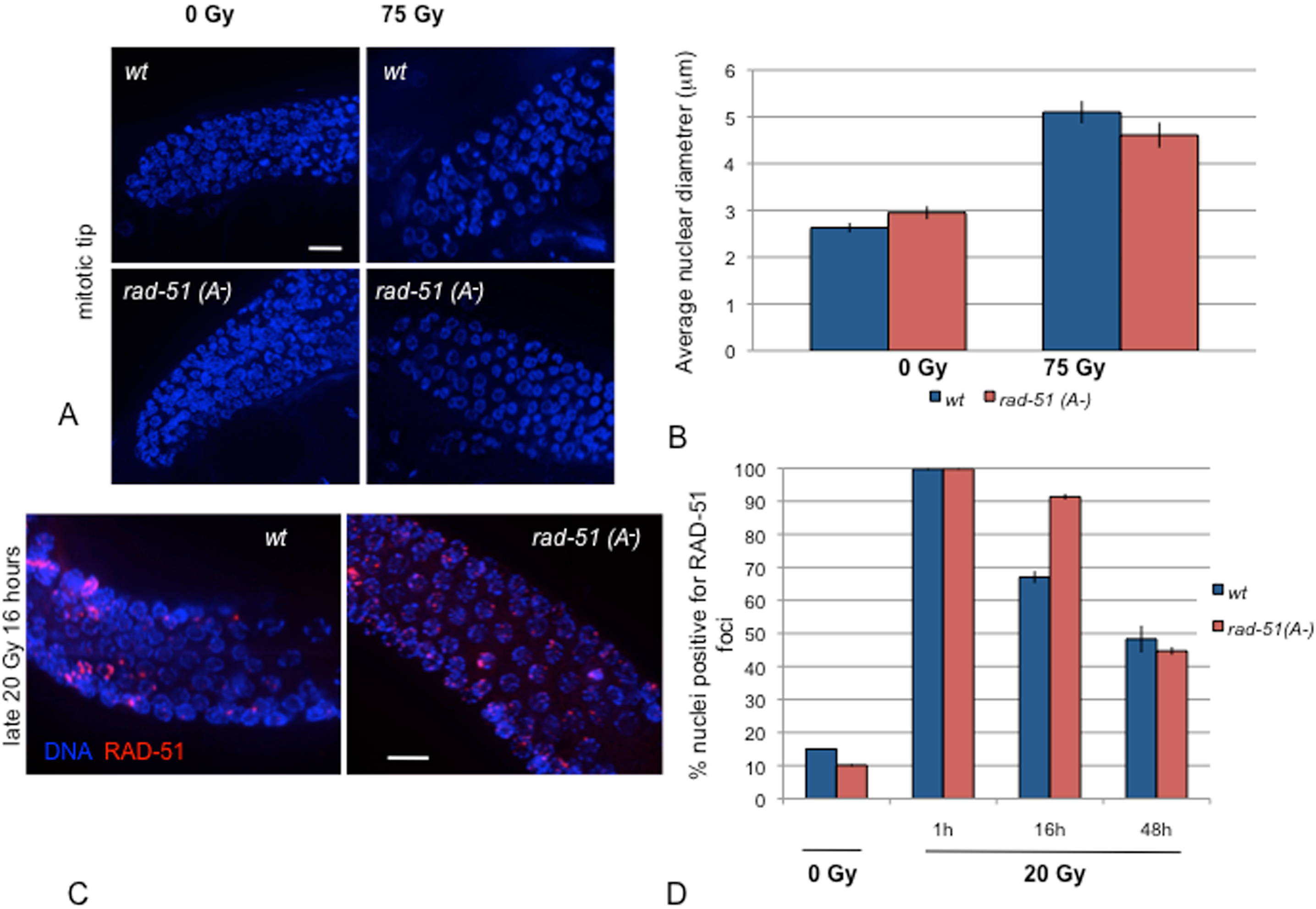
The DNA damage checkpoint is activated in the *rad-51(A^-^)* strain after IR treatment. (A) Representative images of mitotic nuclei before and after treatment with IR (75Gy) in wt and *rad-51(A^-^)* strains. Scale bar, 10 μm (B) Average nuclear diameter of mitotic nuclei before and after treatment with IR (75Gy) in wt and *rad-51(A^-^)* strains. Number of observed nuclei: *wt* 0 Gy: 85, *rad-51(A^-^)* 0 Gy: 130; *wt* 75 Gy: 72, *rad-51(A^-^)* 75 Gy: 106. Mann-Whitney U test: P (*wt* 0 Gy *vs rad-51(A^-^)* 0 Gy) = 0.25, P (*wt* 75 Gy *vs rad-51(A^-^)* 75 Gy) = 0.52, P (*wt* 0 Gy *vs wt* 75 Gy) < 0.0005, P (*rad-51(A^-^)* 0 Gy *vs rad-51(A^-^)* 75 Gy) < 0.0001. (C)) Immunolocalization with anti-RAD-51 in pachytene of wild type and *rad-51(A^-^)* strains 16 h after IR treatment (20Gy). Scale bar, 10 μm (D) Quantification of germ nuclei positive for RAD-51 staining in wild type and rad-51(*A*^-^)strains 1, 16 or 48 hrs after treatment with IR (20Gy). Genotypes are indicated in the color legend on the right side of the chart. Number of observed nuclei: *wt* 0 Gy: 152; *rad-51(A^-^)*0 Gy*: 160; wt 2*0 Gy 1h: 400; *rad-51(A^-^)* 20 Gy 1h*: 560; wt 2*0 Gy 16h: 941; *rad-51(A^-^)* 20 Gy 16h: 705; *wt 2*0 Gy 48h: 584; *rad-51(A^-^)* 20 Gy 48h: 1045 χ ^2^ test: P (*wt* 0 Gy *vs rad-51(A^-^)* 0 Gy) = 0.02, P (*wt 2*0 Gy 1h *vs rad-51(A^-^)* 20 Gy 1h)= 0.6, 3, P (*wt 2*0 Gy 16h *vs rad-51(A^-^)* 20 Gy 16h) < 0.0001, P (*wt 2*0 Gy 48h *vs rad-51(A^-^)* 20 Gy 48h) = 0.18.

Since the DNA damage checkpoint seems to be activated in irradiated *rad-51(A^-^)* gonads, we wondered whether failure in apoptosis execution might be associated with a delay in RAD-51 foci un(?)loading in late pachytene as observed in the *ufd-2* mutant by Ackermann *et al.* 2016. We therefore performed RAD-51 immunolocalization experiments on nematodes that have been irradiated at 20 Gy, and we analysed the kinetics of RAD-51 disassemby on pachytene chromosomes one hour after IR. Both the wild type and the *rad-51 (A^-^)* mutant gonads show about 100% of the nuclei in late pachytene to be positive for RAD-51 foci. However, more than 90% of the late pachytene *rad-51 (A^-^)* mutant nuclei still shows RAD-51 foci 16 hrs after IR unlike what is observed in wild type, where about a third of the pachytene nuclei are devoid of RAD-51 foci. Eventually, 48 hrs after IR, in both strains less than 50% of the pachytene nuclei show detectable foci (Figure 5 C and D).

### RAD-51 isoform A is necessary to induce apoptosis in response to endogenous DNA damages

The *brc-1* mutant shows a permanence of RAD-51 foci in middle to late pachytene thought to represent stalled intersister homologous repair intermediates (Adamo *et al*. 2008 and Figure 6A). We analysed the RAD-51 foci appearance/unloading trend in the double mutant *brc-1;rad-51(A^-^)* and found that the level of RAD-51 foci in the double mutant is higher in the transition zone, but lower in pachytene compared to the *brc-1* mutant control (Figure 6A and Table S2). Furthermore, the *brc-1;rad-51(A^-^)* double mutant shows six DAPI stained bodies at diakinesis similarly to the wildtype, the *rad-51(A^-^)* and the *brc-1* single mutants (Figure 6B).

**Figure 6.**
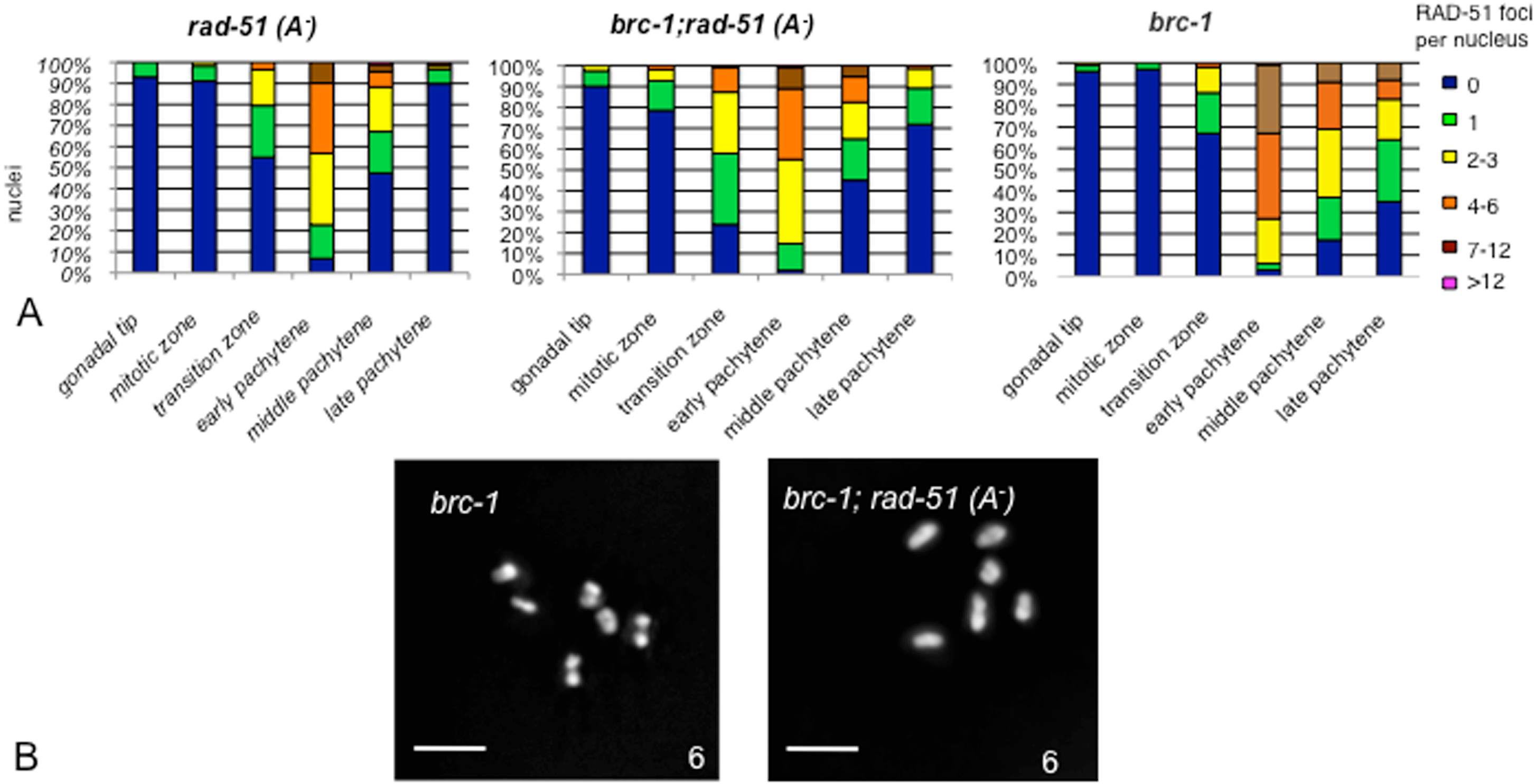
Mutation in *rad-51*(A^-^) does not delay RAD-51 foci downloading in *brc-1*. (A) Quantification of RAD-51 foci in the germlines of the indicated genotypes. An average of 130 nuclei for each gonad region was scored for each getopype. Statistical analyses are reported in table S2. (B) Representative images of diakinesis nuclei of the indicated genotype stained with DAPI. The number of DAPI-stained bodies is indicated at the bottom right of each panel. Scale bar 2 μm.

In absence of the synaptonemal complex, RAD-51 foci are present at high level on chromosomes during middle to late pachytene (Colaiacovo *et al.* 2003). The double mutant *syp-2;rad-51(A^-^)* shows a similar trend to the single mutant *syp-2* (Figure S4).

Lack of the CEP-1 protein suppresses the apoptotic response observed in some meiotic mutants, such as *brc-1* or *syp-2* (Bhalla and Dernburg 2005; Adamo *et al.* 2008; Silva *et al.* 2013). Therefore, we investigated whether DNA damage induced apoptosis in meiotic mutants also requires *rad-51(A^-^).* We observe that *brc-1;rad-51(A^-^)* double mutant displays apoptotic levels similar to the wild-type controls and to the *brc-1;cep-1* double mutant (see Figure 7A and Table S3). Furthermore, the level of apoptosis detected in the *syp-2;rad-51(A^-^)* double mutant is significant lower than that observed in *syp-2* single mutant (Figure 7B and Table S3) and similar to the *syp-2;cep-1* (and to that previously observed in *syp-2;spo-11* and *syp-2 msh-4* double mutants Silva *et al.* 2013), in which just the synapsis checkpoint is activated.

**Figure 7.**
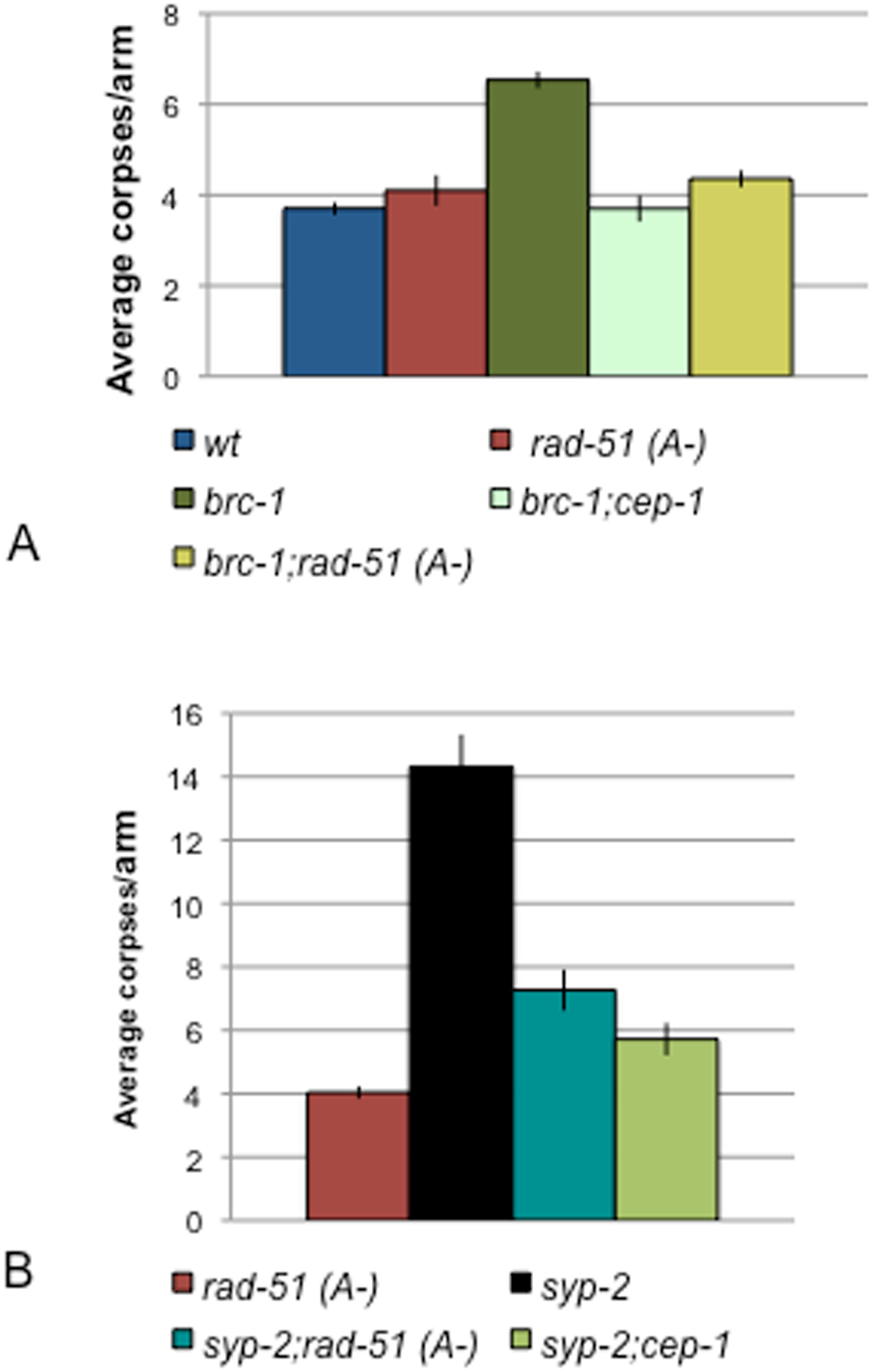
RAD-51 A is necessary to induce apoptosis in DNA repair defective mutants. Average apoptosis levels of the indicated genotypes. The y axis shows the average number of SYTO-12 labeled nuclei per gonadal arm. The genotype are shown in the color legend at the bottom of each chart. Number of observed gonads: *wt*: 71, *rad-51(A^-^)*: 88, brc-1: 61, *brc-1;rad-51(A^-^):* 72, *brc-1;cep-1:* 94, *syp-2:* 30, *syp-2;rad-51(A^-^):* 52*;syp-2;cep-1:* 39. Error bars correspon to standard error of the means (S.E.M.) calculated from at least three independent experiments. Student’s *t*-tests are shown in table S3.

Therefore we can conclude that the RAD-51 isoform A is necessary for DNA damage dependent apoptosis in response to endogenous DNA damages.

## Discussion

The recA-like protein RAD-51 is known to catalyze the strand transfer between homologous chromosomes or sister chromatids during meiosis and DNA repair. In most eukaryotes another recA like protein is also involved in this mechanism during meiosis. Such a protein is however missing in *C. elegans*. The *C. elegans rad-51* gene is transcribed into three mRNAs potentially coding for alternative protein isoforms. In the course of our experiments we demonstrated that the RAD-51 longest isoform performs a different function from the shorter forms. In absence of the RAD-51 long isoform A, in fact, we observe correct chromosome segregation as no difference is observed in male frequency or embryonic lethality between *rad-51(A^-^)* mutant and the wild type (Figure 1 and Table 1). The only abnormal phenotype observed is larval arrest. Defects in germline proliferation and vulva development are significantly enhanced after gamma rays treatment. This observation suggests an involvement of the RAD-51 long isoform A in genome stability during larval development.

Petermann *et al.* 2010 have suggested that in mammalian cells restart of stalled replication forks and HR repair of collapsed replication forks require two distinct RAD51-mediated pathways. They demonstrated that stalled replication forks are efficiently restarted in a RAD51-dependent process that does not trigger homologous recombination (HR) suggesting that RAD51-mediated strand invasion supports fork restart while RAD51-dependent HR is triggered only in case of collapsed replication forks, without apparent restart. It is possible that RAD-51 isoform A may be involved during late development either in DNA repair or in restarting blocked replication forks.

The frequency and distribution of RAD-51 foci on chromosomes along the Meiosis I prophase in *rad-51(A^-^)* are indistinguishable from wild type. This seems to suggest that the isoform A is not preferentially loaded at the sites of SPO-11 induced DSBs (Figure 1B and Table S2).

In meiosis, the repair of DSBs by HR leads to the formation of COs and non crossovers(NCOs); during *C. elegans* oogenesis, only half of DSBs become COs: each chromosome pair has one (obligatory) CO which is required to lock the homologues together during the first meiotic division resulting in proper chromosome segregation. All the others DSBs, once the COs are produced, will be repaired using the sister chromatid as template. We have investigated on the possible involvement of the RAD-51 long isoform A during SPO-11 induced DSBs repair either in a genetic background in which the synaptonemal complex is missing (and therefore the sister chromatid is the only available template for HR) or in a genetic background where homologous DNA repair on the sister chromatid is defective. In either double mutant *syp-2;rad-51(A^-^)* and *brc-1;rad-51(A*^-^) the average number of diakinesis DAPI stained bodies are indistinguishable from those observed in the *syp-2* and *brc-1* single mutants. These observations furtherl support our hypothesis that the RAD-51 long isoform A is not involved in DNA repair during meiosis I.

RAD-51 long isoform A appears, instead, to have a fundamental role in inducing apoptosis following DNA damage. Following gamma rays treatment in *rad-51(A*^-^) mutant, as well as in wild type, the DNA damage checkpoint is activated (Figure 5A), and a large number of RAD-51 foci are loaded on chromosomes along the entire gonad (figure 3). Nevertheless, the level of germline apoptosis remains at basal levels as in *rad-51(A^-^)* or wildtype untreated hermaphrodites. These data apparently contradict what has been observed so far in the *rad-51(lg8701)* deletion mutant, where none of the RAD-51 isoforms are expressed (Alpi *et al.* 2003, Silva *et al.* 2013) and the level of apoptosis is at very high levels. However the *rad-51(lg8701)* deletion mutant produce 100% dead embryos and therefore only the first homozygous generation can be observed. We observed, by immunolocalization, that a small but significant amount of RAD-51 foci, likely of maternal origin, persists in the germline of the *rad-51(lg8701)* homozygous progeny of the *rad-51(lg8701) IV/nT1 [let-?(m435)] (IV;V)* balanced heterozygous hermaphrodites. Although the small amount of RAD-51 short isoforms is not sufficient to repair the SPO-11 induced DSBs, it is likely that the amount of RAD-51 long isoform A of maternal origin is sufficient to induce germline apoptosis. In support of such hypothesis we observed that the level of germline apoptosis drops to wild type levels in the *rad-51(lg8701)* homozygous progeny of a *rad-51(lg8701); rad-51(A^-^)* heterozygous hermaphrodites (figure 4C and S3) where the RAD-51 long isoform A has been absent for two generations.

In *rad-51(A^-^)* as in *ufd-2* mutants RAD-51 foci have a delay in the kinetics of disassembly after gamma ray treatment. It has been hypothesized that the RAD-51 disassembly itself may trigger the induction of apoptosis (Ackerman *et al.* 2016). However, we cannot exclude that the delay in RAD-51 disassembly may be a consequence and not the cause of apoptosis induction. In fact, all the apoptotic corpses, not only those due to DNA damage, but even those due to physiological apoptosis, always appear overloaded with RAD-51 foci. Furthermore, in many meiotic mutant RAD-51 foci accumulate in late pachytene. They persist, for example, both in msh-4/5 mutants, where DNA damage dependent apoptosis is blocked, and in *brc-1* and *syp-2* mutants where high level of damage induced apoptosis is present.

The RAD-51 long isoform A is, in fact, also involved in inducing apoptosis in response to endogenously induced DNA damage such as in DNA repair defective mutants. We previously demonstrated that in the *brc-1* mutant DNA repair on the sister chromatid is defective in meiosis, that a high level of HR intermediates accumulate in late pachytene, and that DNA damage apoptosis is induced (Adamo *et al.* 2008). In the *brc-1;rad-51(A^-^)* double mutant the level of apoptotic corpses drops to the physiological wild type level and RAD-51 foci (representing HR intermediates) do not accumulate in late pachytene chromosomes as they do in the *brc-1* single mutant. Therefore, in absence of *rad-51(A^-^),* DNA damage dependent apoptosis is not induced in a *brc-1* background.

In the *syp-2* mutant, the synaptonemal complex is not present and COs are not formed. Although SPO-11 induced DSBs will be eventually repaired using the sister chromatid as template, RAD-51 foci accumulate on late pachytene chromosomes (Colaiacovo *et al.* 2003) and both the pairing checkpoint and the DNA repair checkpoint are activated with a significant increase in germline apoptosis (Bhalla and Dernburg 2005). Also in this genetic background, *cep-1* dependent apoptosis disappears in absence of the RAD-51 long isoform A and only the physiological and the pairing checkpoint dependent corpses persist.

We conclude that the RAD-51 long isoform A has a key role in the induction of apoptosis downstream of the activation of the DNA damage checkpoint. Although DNA damage and repair defects can be present along the whole gonad, apoptotis is allowed to start only just before diplotene stage. Therefore, a complex pattern of interactions seems to be active. Further studies will be required in *C. elegans* to understand the cross-talking among the genes involved in the meiotic apoptotic induction including, beside *rad-51,* also *msh-4/5, zhp-3* (Silva *et al.* 2013), and *ufd-2* (Ackermann *et al.* 2016) and to identify the specific physical interactions and steps of the apoptotic response at which each gene is required.

## Figure legends

**Figure S1 *rad-51(A^-^)* mutant**

*rad-51(A^-^)* is a 56-bp deletion/79-bp insertion obtained by CRISP/Cas9 mediated genome editing. The deletion includes 45-bp of the 5’ UTR, the first ATG and part of the first exon.

(A) Schematic representation of the *rad-51* gene in the 5’ region up to the end of the third exon. Boxes and straight lines represent exons and introns, respectively.

(B) Deletion and insertion sequence are show in Italic. The first exon is indicated in uppercase.

(C) N-terminal portion of predicted isoforms in *wt*. In *rad-51(A^-^)* the isoform A is absent.

**Figure S2 RAD-51 A in the larval development**

(A) Quantification of larval arrests and egg laying defects in *wt* and *rad-51(A^-^)* mutant. At the top untreated nematodes are shown, at the bottom nematodes that have been irradiated at the stage L1/L2 at 120 Gy and screened 52 hrs later.

χ ^2^ test: P (*wt* 0 Gy *vs rad-(A^-^)* 0 Gy) = 0.0085, P (*wt 12*0 Gy *vs rad-51(A^-^)* 120 Gy) < 0.0001, P (*wt* 0 Gy *vs wt* 120 Gy) < 0.0001, P (*vs rad-51(A^-^)* 0 Gy *vs vs rad-51(A^-^)* 120 Gy) < 0.0001.

(B) Quantification of vulvaless phenotypes in animals irradiated at 120 Gy at the stage L1/L2 and screened 52 hrs later. Genotypes are indicated in the color legend at the bottom of the chart.

The y axis shows the percent of the adults with vulvaless phenotypes.

**Figure S3 genetic cross**

P_0_ *rad-51(A^-^)* homozygous hermaphrodites were crossed with P_0_ *rad-51(lg8701) dpy-13 /nT1 [unc-? (n754) let-? qls50](IV;V)* males. The heterozygous *rad-51(A^-^)/rad-51(lg8701) dpy-13* hermaphrodite F1 was selected, isolated and left to segregate. In the F2 the 25% of the progeny that was homozygous for *rad-51(lg8701) dpy-13* was selected and analyzed.

**Figure S4 Immunostaining of RAD-51 in *syp-2; rad-51(A^-^)* strains**

Representative images of middle pachytene nuclei in *syp-2* and *syp-2; rad-51(A^-^)*immunostained for RAD-51.

Scale bar, 10 μm.

## Acknowledgements

We thank Adriana La Volpe, John Pulitzer for useful discussions and critical reading of the paper. Some nematode strains used in this study were provided by the *Caenorhabditis* Genetics Center, which is funded by the NIH National Center for Research Resources (NCRR). We thank Monica Colaiacovo for providing a strain. This study was started as part of the Doctoral thesis of Marcello Germoglio in Biomolecular Science, at the Second University of Naples, who was the recipient of a predoctoral fellowship from Telethon Grant GGP11076.

## References

Ackermann, L., M. Schell, W. Pokrzywa, É. Kevei, A. Gartner, et al., 2016 E4 ligase-specific ubiquitination hubs coordinate DNA double-strand-break repair and apoptosis. Nat Struct Mol Biol. 23(11):995–1002.

Adamo, A., S.J. Collis, C.A. Adelman, N. Silva, Z. Horejsi, et al., 2010 Preventing nonhomologous end joining suppresses DNA repair defects of Fanconi anemia. Mol Cell. 39(1):25–35.

Adamo, A., P. Montemauri, N. Silva, J.D. Ward, S.J. Boulton, et al., 2008 BRC-1 acts in the inter-sister pathway of meiotic double-strand break repair. EMBO Rep. 9(3):287–92.

Adamo, A., A. Woglar, N. Silva, A. Penkner, V. Jantsch, et al., 2012 Transgene-mediated cosuppression and RNA interference enhance germ-line apoptosis in *Caenorhabditis elegans*. Proc Natl Acad Sci U S A. 109(9):3440–5.

Alpi, A., P. Pasierbek, A. Gartner, and J. Loidl, 2003 Genetic and cytological characterization of the recombination protein RAD-51 in *Caenorhabditis elegans*. Chromosoma. 112(1):6–16.

Bhalla, N., and A.F. Dernburg, 2005 A conserved checkpoint monitors meiotic chromosome synapsis in *Caenorhabditis elegans*. Science. 310(5754):1683–6.

Bishop, D.K., D. Park, L. Xu, and N. Kleckner, 1992 DMC1: a meiosis-specific yeast homolog of *E. coli* recA required for recombination, synaptonemal complex formation, and cell cycle progression. Cell. 69(3):439–56.

Brenner S., 1974 The genetics of *Caenorhabditis elegans*. Genetics. 77:71–94

Brown, M.S., and D.K. Bishop, 2014 DNA strand exchange and RecA homologs in meiosis. Cold Spring Harb Perspect Biol. 7(1):a016659.

Colaiacovo, M.P., 2006 The many facets of SC function during *C. elegans* meiosis. Chromosoma. 115: 195–211.

Colaiácovo, M.P., A.J. MacQueen, E. Martinez-Perez, K. McDonald, A. Adamo, et al., 2003 Synaptonemal complex assembly in *C. elegans* is dispensable for loading strand-exchange proteins but critical for proper completion of recombination. Dev Cell. 5(3):463–74.

Couteau, F., and M. Zetka, 2005 HTP-1 coordinates synaptonemal complex assembly with homolog alignment during meiosis in *C. elegans*. Genes Dev. 19, 2744–2756.

Dickinson, D.J., Ward, J.D., Reiner, D.J. and Goldstein B 2013 Engineering the *Caenorhabditis elegans* genome using Cas9-triggered homologous recombination. Nat. Methods. 10:1028–1034

Gartner, A., S. Milstein, S. Ahmed, J. Hodgkin, and M.O. Hengartner, 2000 A conserved checkpoint pathway mediates DNA damage--induced apoptosis and cell cycle arrest in *C. elegans*. Mol Cell. 5(3):435–43.

Gumienny, T.L., E. Lambie, E. Hartwieg, H.R. Horvitz, and M.O. Hengartner, 1999 Genetic control of programmed cell death in the *Caenorhabditis elegans* hermaphrodite germline. Development. 126(5):1011–22

Hillers, K.J., V. Jantsch, E. Martinez-Perez, and J.L. Yanowitz, 2017 Meiosis. WormBook. 4:1–43.

Hofmann, E.R., S. Milstein, S.J. Boulton, M. Ye, J.J. Hofmann, et al., 2002 *Caenorhabditis elegans* HUS-1 is a DNA damage checkpoint protein required for genome stability and EGL-1-mediated apoptosis. Curr Biol. 12(22):1908–18.

Jantsch V, P. Pasierbek, MM. Mueller, D. Schweizer, M. Jantsch, et al., 2004 Targeted gene knockout reveals a role in meiotic recombination for ZHP-3, a Zip3-related protein in Caenorhabditis elegans Mol Cell Biol. Sep;24(18):7998–8006.

Kelly, K.O., A.F. Dernburg, G.M. Stanfield, and A.M. Villeneuve, 2000 *Caenorhabditis elegans msh-5* is required for both normal and radiation-induced meiotic crossing over but not for completion of meiosis. Genetics. 156: 617–630.

Kim, H.M., and M.P. Colaiácovo, 2014 ZTF-8 interacts with the 9-1-1 complex and is required for DNA damage response and double-strand break repair in the *C. elegans* germline. PLoS Genet. 10(10):e1004723.

Kim, H.M., and M.P. Colaiácovo, 2015 New Insights into the Post-Translational Regulation of DNA Damage Response and Double-Strand Break Repair in *Caenorhabditis elegans*. Genetics. 200(2):495–504.

Kleckner, N., 1996 Meiosis: how could it work? Proc Natl Acad Sci U S A. 93(16):8167–74.

La Volpe, A., and M. Barchi, 2012 Meiotic double strand breaks repair in sexually reproducing eukaryotes: we are not all equal. Exp Cell Res. 318(12):1333–9.

Machovina, T.S., R. Mainpal, A. Daryabeigi, O. McGovern, D. Paouneskou, et al., 2016 A Surveillance System Ensures Crossover Formation in *C. elegans*. Curr Biol. 26(21):2873–2884.

Meneely, P.M., O.L. McGovern, F.I. Heinis, and J.L. Yanowitz, 2012 Crossover distribution and frequency are regulated by *him-5* in *Caenorhabditis elegans*. Genetics. 190(4):1251–66.

Mets, D.G., and B.J. Meyer, 2009 Condensins regulate meiotic DNA break distribution, thus crossover frequency, by controlling chromosome structure. Cell. 139(1):73–86.

Moser, S.C., S. Von Elsner, I. Bussing, A. Alpi, R. Schnabel et al. 2009 Functional dissection of Caenorhabditis elegans CLK-2/TEL2 cell cycle defects during embryogenesis and germline development. PLoS Genet; 5: e1000451.

Penkner, A., L. Tang, M. Novatchkova, M. Ladurner, A. Fridkin, et al., 2007 The nuclear envelope protein Matefin/SUN-1 is required for homologous pairing in *C. elegans* meiosis. Dev Cell. 12(6):873–85.

Petermann, E., M.L. Orta, N. Issaeva, N. Schultz, and T. Helleday, 2010 Hydroxyurea-stalled replication forks become progressively inactivated and require two different RAD51-mediated pathways for restart and repair Mol Cell. Feb 26;37(4):492–502.

Rinaldo, C., P. Bazzicalupo, S. Ederle, M. Hilliard, and A. La Volpe, 2002 Roles for *Caenorhabditis elegans rad-51* in meiosis and in resistance to ionizing radiation during development. Genetics. 160(2):471–9.

Rinaldo, C., S. Ederle, V. Rocco, and A. La Volpe, 1998 The *Caenorhabditis elegans* RAD51 homolog is transcribed into two alternative mRNAs potentially encoding proteins of different sizes. Mol Gen Genet. 260(2-3):289–94.

Rosu, S., D.E. Libuda, and A.M. Villeneuve, 2011 Robust crossover assurance and regulated interhomolog access maintain meiotic crossover number. Science. 334(6060):1286–9.

Rutkowski, R., R. Dickinson, G. Stewart, A. Craig, M. Schimpl, et al., 2011 Regulation of *Caenorhabditis elegans* p53/CEP-1-dependent germ cell apoptosis by Ras/MAPK signaling. PLoS Genet. 7(8):e1002238.

Schumacher, B., M. Hanazawa, M.H. Lee, S. Nayak, K. Volkmann, E.R. Hofmann, M. Hengartner, T. Schedl, and A. Gartner, 2005 Translational repression of *C. elegans* p53 by GLD-1 regulates DNA damage-induced apoptosis. Cell. 120(3):357–68.

Schumacher, B., K. Hofmann, S. Boulton, and A. Gartner, 2001 The *C. elegans* homolog of the p53 tumor suppressor is required for DNA damage-induced apoptosis. Curr Biol. 11: 1722–1727.

Silva, N., A. Adamo, P. Santonicola, E. Martinez-Perez, and A. La Volpe, 2013 Pro-crossover factors regulate damage-dependent apoptosis in the *Caenorhabditis elegans* germ line. Cell Death Differ. 20(9):1209–18.

Zalevsky, J., A.J. MacQueen, J.B. Duffy, K.J. Kemphues, and A.M. Villeneuve, 1999 Crossing over during *Caenorhabditis elegans* meiosis requires a conserved MutS-based pathway that is partially dispensable in budding yeast. Genetics. 153: 1271–1283.

